# Machine learning reveals time-varying microbial predictors with complex effects on glucose regulation

**DOI:** 10.1101/2020.08.13.250423

**Authors:** Oliver Aasmets, Kreete Lüll, Jennifer M. Lang, Calvin Pan, Johanna Kuusisto, Krista Fischer, Markku Laakso, Aldons J. Lusis, Elin Org

## Abstract

The incidence of type 2 diabetes (T2D) has been increasing globally and a growing body of evidence links type 2 diabetes with altered microbiota composition. Type 2 diabetes is preceded by a long pre-diabetic state characterized by changes in various metabolic parameters. We tested whether the gut microbiome could have predictive potential for T2D development during the healthy and pre-diabetic disease stages. We used prospective data of 608 well-phenotyped Finnish men collected from the population-based Metabolic Syndrome In Men (METSIM) study to build machine learning models for predicting continuous glucose and insulin measures in a shorter (1.5 year) and longer (4.5 year) period. Our results show that the inclusion of gut microbiome improves prediction accuracy for modelling T2D associated parameters such as glycosylated hemoglobin and insulin measures. We identified novel microbial biomarkers and described their effects on the predictions using interpretable machine learning techniques, which revealed complex linear and non-linear associations. Additionally, the modelling strategy carried out allowed us to compare the stability of model performances and biomarker selection, also revealing differences in short-term and long-term predictions. The identified microbiome biomarkers provide a predictive measure for various metabolic traits related to T2D, thus providing an additional parameter for personal risk assessment. Our work also highlights the need for robust modelling strategies and the value of interpretable machine learning.

**Importance:** Recent studies have shown a clear link between gut microbiota and type 2 diabetes. However, current results are based on cross-sectional studies that aim to determine the microbial dysbiosis when the disease is already prevalent. In order to consider microbiome as a factor in disease risk assessment, prospective studies are needed. Our study is the first study that assesses the gut microbiome as a predictive measure for several type 2 diabetes associated parameters in a longitudinal study setting. Our results revealed a number of novel microbial biomarkers that can improve the prediction accuracy for continuous insulin measures and glycosylated hemoglobin levels. These results make the prospect of using microbiome in personalized medicine promising.

## Background

The prevalence of type 2 diabetes (T2D) has more than doubled since 1980, resulting in a huge burden on the health care system worldwide (1). In order to fight the epidemic of T2D and improve public health, understanding of the first stages of this disease is necessary for preventive actions. Recently, the bacterial communities residing in our intestines have become a topic of interest as a potential way to prevent the development of glucose dysregulation. The microbiome has been shown to modulate a variety of physiological functions, such as gut permeability, inflammation, glucose metabolism and fatty acid oxidation, supporting an important role of the microbiome in the pathophysiology of T2D (2).

Numerous studies have already reported changes in the gut microbiome in subjects with T2D or prediabetes compared to healthy individuals (3–5). Although there is information that the abundance of bacteria such as *Roseburia* and *Bifidobacteria* is altered in subjects with T2D (2), compelling evidence that supports the use of gut microbiome as a predictive tool for T2D is lacking, as a majority of the findings are based on cross-sectional studies. However, in order to assess the microbiome as a prognostic tool for T2D, prospective studies are needed.

T2D is a heterogeneous disease with multiple pathophysiological pathways involved (6). Thus, in order to fully understand the role of microbiome in the risk of T2D, a case-control design might not be sufficient. As the progression of the disease is a continuous process, detailed data about metabolic outcomes such as continuous glucose and insulin measurements could help to unravel the disease mechanisms involving the microbiome.

Together with heterogeneity in the first stages T2D, the gut microbiome itself is known to be highly personalized (7, 8). Variability in continuous metabolic outcomes and gut microbiome lead to difficulties in reproducing the results obtained and raises the need for robust modelling strategies. Machine learning methods have been shown to capture various complex association patterns from different data types. Although machine learning has become popular in microbiome studies as well, the ability of the algorithms to provide robust results remains unclear (9, 10).

We now report the application of a random forest algorithm on microbiome data to predict multiple continuous metabolic outcomes that influence the development of T2D in a longitudinal study setting. We identify microbial biomarkers for the metabolic outcomes and describe their effects on the predictions using interpretable machine learning techniques. In addition, we show that there are significant differences in the identified biomarkers between long- and short follow-up periods. We also show how the modelling procedure significantly influences the results.

## Results

### Study design

We used prospective data of well phenotyped Finnish men collected from a population-based Metabolic Syndrome In Men (METSIM) study. A comprehensive machine learning strategy was implemented to identify microbial biomarkers and their effect on numerous metabolic traits. Graphical overview of the study design and modelling procedure is shown in **Figure 1**. Random forest models were trained to predict the metabolic outcomes of interest in the follow-up using baseline microbiome (MB), metabolic outcomes (MO) and additional covariates (CoV) such as body mass index and age as predictors. To evaluate the effect of microbiome, models including microbial predictors were compared to models excluding microbial predictors. In order to assess the temporal changes in biomarker selection and predictive performance, independent prospective models were trained for the 18-month and 48-month follow-up period. To evaluate the model generalizability and stability, model training was repeated 200 times with different train-test split made each run. Permutation feature importance metrics were used to identify microbial biomarkers. Finally, accumulated local effects methodology was used to plot the effect of the microbial biomarkers for predicting the corresponding metabolic trait.

**Figure 1.**
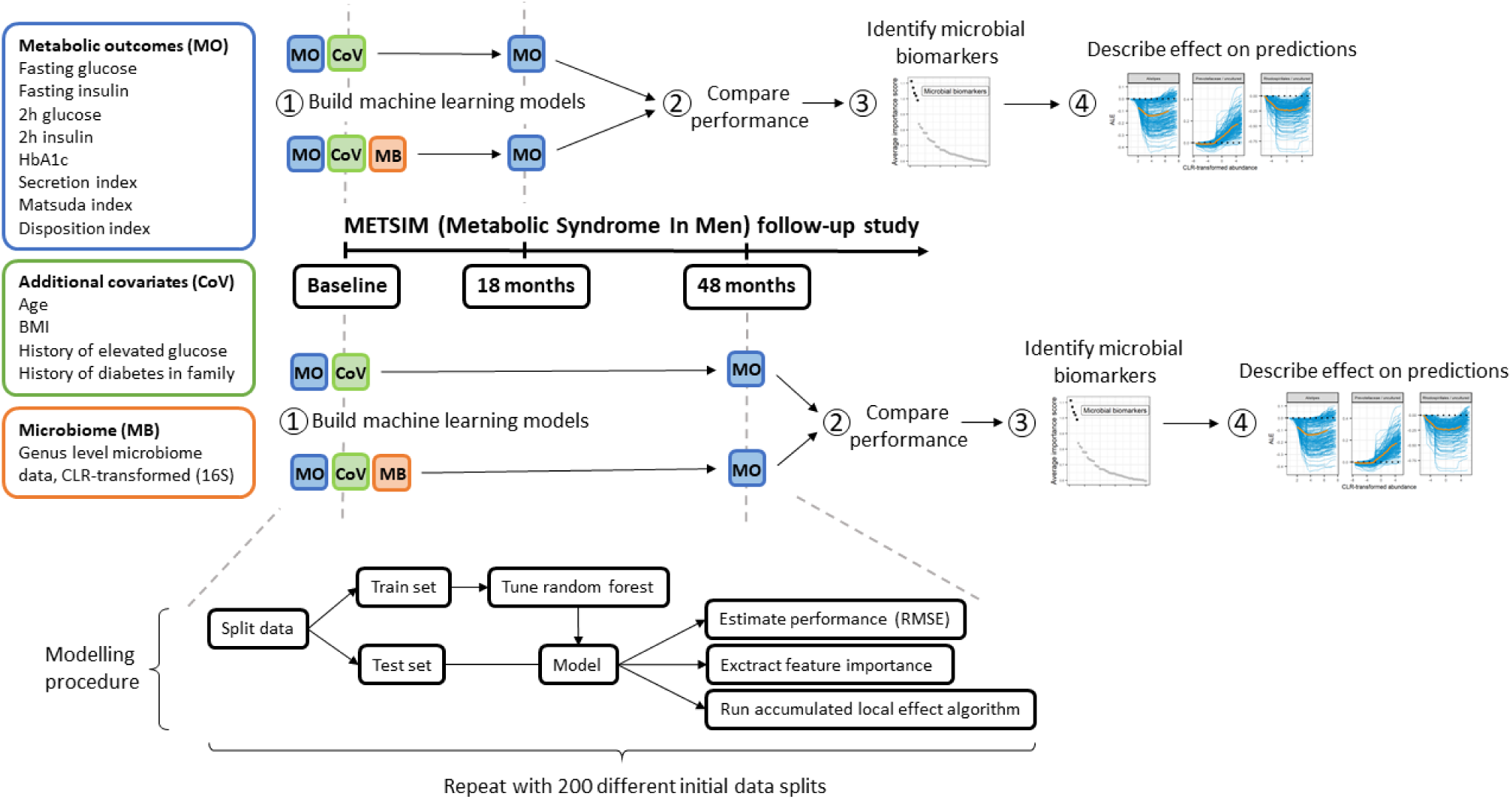
Study design and modelling procedure.

### Model stability and generalizability

In the first step we tested whether we could improve the prediction of metabolic outcomes using microbiome data as an additional predictor. Human gut microbiome is known to be highly variable and personalized (7, 8). Thus, estimating the robustness of the predictive models is essential. The problem with microbiome data based on our experience is that the performance of the model might be highly dependent on the initial data split to training and test sets. The models were run 200 times with different initial splits to assess the impact of the data split. **Table 1** summarizes the obtained results.

**Table 1.** Model stability and generalizability.

| Trait | 18-month time frame |  | 48-month time frame |  |
| --- | --- | --- | --- | --- |
|  | Mean (sd)<br>difference in<br>RMSE | # models<br>including<br>microbiome<br>performing better | Mean (sd)<br>difference in RMSE | # models including<br>microbiome<br>performing better |
| Fasting glucose | 0.001 (0.0594) | 99 (49.5%) | -0.006 (0.0641) | 112 (56%) |
| 2h glucose | -0.02 (0.217) | 118 (59%) | 0.07 (0.332) | 73 (36.5%) |
| Fasting insulin | 0.20 (1.04) | 73 (36.5%) | -0.29 (1.080) | 137 (68.5%) * |
| 2h insulin | -3.23 (10.840) | 141 (70.5%) * | -1.42 (12.304) | 122 (61%) * |
| HbA1c | -0.005 (0.0305) | 129 (64.5%) * | -0.002 (0.0360) | 111 (55.5%) |
| Secretion index | -0.36 (4.949) | 122 (61%) * | -0.77 (3.254) | 138 (69%) * |
| Matsuda index | 0.07 (0.573) | 90 (45%) | -0.01 (0.569) | 103 (51.5%) |
| Disposition index | 4.42 (26.590) | 77 (38.5%) | 2.01 (16.251) | 86 (43%) |

Mean differences in root-mean-square error (RMSE) between models including microbial predictors and models excluding microbial predictors. Negative value indicates a model including microbial predictors outperforming the model excluding microbial predictors. \* shows statistically significant results according to the binomial test after Bonferroni correction.

These results highlight the variability in performance estimates occurring due to the data split. Out of 200 data splits, the number of models that took advantage of using microbial predictors varies around 100 which implies that the data split plays an important role in the outcome. Our results suggest that for the 18-month time frame, microbiome as a predictor can improve the prediction accuracy for secretion index, glycosylated hemoglobin (HbA1c) and 2h insulin levels. For secretion index, models including microbial predictors outperformed simpler models in 61% of the cases, for 2h insulin in 70.5% of the cases and for HbA1c in 64.5% of the cases. For a 48-month time frame the microbiome improves the prediction model for the secretion index, fasting insulin and 2h insulin. For secretion index, models including microbial predictors outperformed simpler models in 69% of the cases, for 2h insulin in 61% of the cases, and for fasting insulin in 68.5% of the cases.

Remarkably, the variation in differences in root-mean-square error (RMSE) between the model including microbial predictors and model excluding microbial predictors over the 200 runs is large. Due to the high variability, the level of improvement in prediction accuracy when microbiome data are used remains unclear.

### Novel predictive microbial biomarkers for metabolic outcomes

In order to find microbial markers that are predictive for the metabolic outcomes, average feature importance scores over 200 runs were compared. **Figure 2** shows the average importance score of top 50 microbial predictors for metabolic outcomes that took advantage of using microbial predictors. It can be seen that certain microbial predictors significantly stand out for each metabolic outcome and time frame combination.

**Figure 2.**
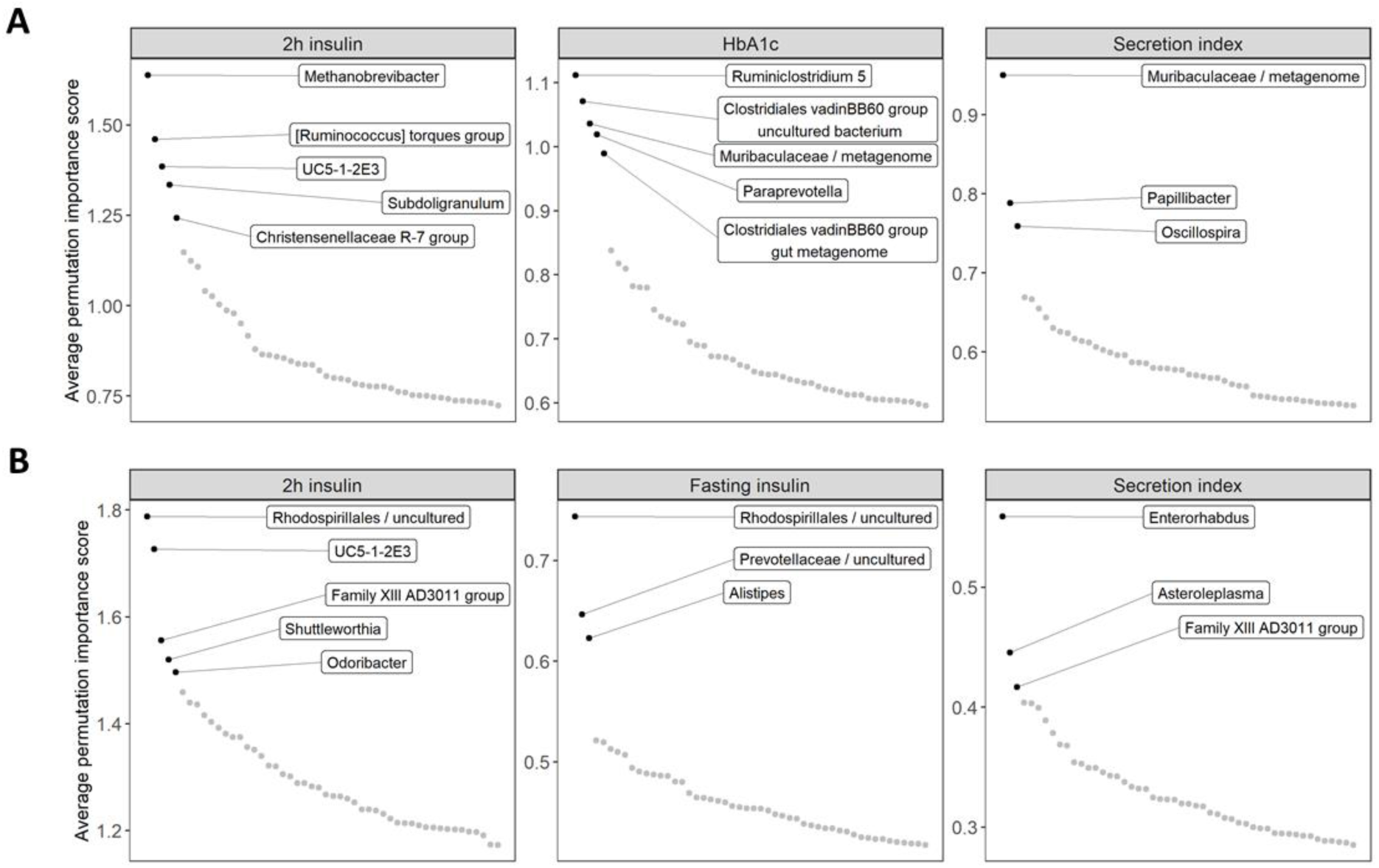
Average feature importance scores for top 50 microbial markers. Highlighted taxa are considered the most significant biomarkers. (**A**) Predictors for 18-month follow-up. (**B**) Predictors for 48-month follow-up.

For a 18-month time frame (**Figure 2A, Supplementary Table S2**), the most important microbial predictors for 2h insulin include genus *Methanobrevibacter* and numerous genera from phylum *Firmicutes* such as *[Ruminococcus] torques group, UC5-1-2E3, Subdoligranulum* and *Christensenellaceae R-7 group*. Predictors for HbA1c are genus *Ruminiclostridium 5*, genus *Paraprevotella*, unclassified member of family *Muribaculaceae* and members of *Clostridiales vadinBB60 group*. Unclassified member of the family *Muribaculaceae* together with *Papillibacter* and *Oscillospira* are significant predictors for secretion index.

For the 48-month time frame (**Figure 2B, Supplementary Table S2**), top predictors for 2h insulin include uncultured *Rhodospirillales* and *UC5-1-2E3*. Distinguishable genera according to the average importance score are also *Family XIII AD3011 group, Shuttleworthia* and *Odoribacter*. Significant predictors for fasting insulin are uncultured *Rhodospirillales*, uncultured *Prevotellaceae* and genus *Alistipes*. For secretion index, genus *Enterohabdus* together with *Asteroleplasma* prove to be the most important predictors, with *Family XIII AD3011 group* slightly standing out.

There is overlap in the most important microbial markers found for predicting different metabolic outcomes. In the 18-month follow-up period, unclassified *Muribaculaceae* is a significant predictor for secretion index and HbA1c. For 48-month follow-up period, *Family XIII AD3011 group* is a predictor for secretion index and 2h insulin and uncultured *Rhodospirillales* is an important predictor for fasting insulin and 2h insulin. Additional overlap can be seen among top 10 microbial predictors according to average permutation importance score (**Supplementary Tables S2 and S3**).

### Interpreting the effect of microbial biomarkers on the predictions

Together with finding the relevant biomarkers, understanding how they influence the predictions is necessary. This task is complicated for most of the machine learning algorithms, which is why they are considered “gray-box” or “black-box” methods. Recently, much attention has been put into explaining the predictions of such models. Here, we implemented accumulated local effect (ALE) plots that aim to describe the effect of a certain predictor on the metabolic outcome independently of the remaining predictors (11). Accumulated local effect plots for previously highlighted most significant microbial biomarkers are shown in **Figure 3**. Accumulated local effect plots for top 10 microbial predictors are shown in **Supplementary Figures 1 and 2**. In most cases, ALE plots show nonlinear associations between a microbial predictor and metabolic outcome of interest. Although large variability in the effect estimates between the different data-splits can be seen, the shape of the effect stays relatively stable for all microbial predictors.

**Figure 3.**
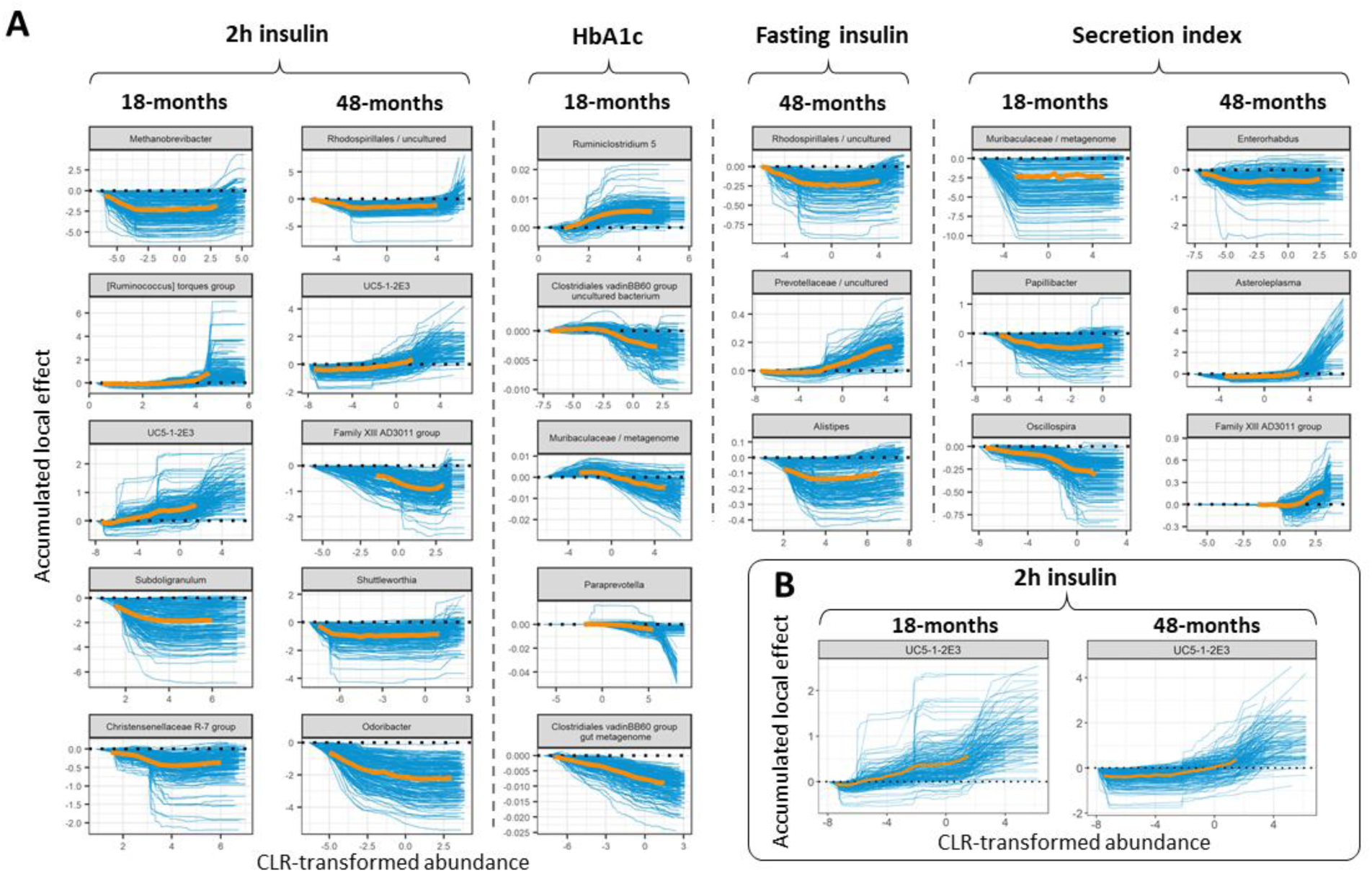
Accumulated local effect (ALE) plots. (**A)** ALE plots for the found microbial biomarkers. (**B**) ALE plots for genus *UC5-1-2E3* found to predict 2h insulin in an 18-month and 48-month follow-up. Blue lines represent effects for each run out of 200, orange lines represent aggregated effects. Aggregated effect is displayed between the 2.5% and 97.5% quantiles of CLR-transformed abundance for the corresponding microbial marker.

Considering the 18-month time frame (**Figure 3A, Supplementary Figure 1**), higher CLR-transformed abundances of genera from the *Lachnospiraceae* family - *[Ruminococcus] torques group* and *UC5-1-2E3* lead to higher predictions for 2h insulin. High CLR-transformed abundances of genera *Subdoligranulum, Methanobrevibacter* and *Christensenellaceae R-7 group* lower the predictions for 2h insulin. For HbA1c, higher CLR-transformed abundance of *Ruminiclostridium 5* leads to higher predictions. On the contrary, high CLR-transformed abundances of bacteria from family *Muribaculaceae*, members of *Clostridiales vadinBB60 group* and *Paraprevotella* reduce the levels of HbA1c. For secretion index, the prediction might depend on the presence-absence of the unclassified genus from family *Muribaculaceae*, because the ALE plot stays relatively stable after an initial decrease from the minimum values of CLR-transformed abundances. High CLR-transformed abundances of *Oscillospira* and *Papillibacter* decrease the predictions for secretion index.

Considering 48-month follow-up period (**Figure 3A, Supplementary Figure 2**), high CLR-transformed abundances of genera *Firmicutes Family XIII AD3011 group, Odoribacter* and unclassified *Rhodospirillales* lead to lower predictions for 2h insulin. In contrast, extremely high CLR-transformed values of genus *UC5-1-2E3* lead to higher predictions. *Shuttleworthia* seems to show presence-absence effect as the drop from the lowest CLR-transformed values lowers the predictions for 2h insulin. For fasting insulin, higher CLR-transformed abundances of unclassified *Rhodospirillales* and *Alistipes* lower the predictions. In contrast, high CLR-transformed abundances of unclassified genus from *Prevotellaceae* family leads to higher predictions for fasting insulin. Interestingly, extremely low values of *Alistipes* lead to higher predictions for fasting insulin compared to when *Alistipes* levels are within 2.5% and 97.5% quantiles. Similar effect for genus *Asteroleplasma* on secretion index can be seen as extremely high CLR-transformed abundance of *Asteroleplasma* leads to drastically higher predictions. Genus *Enterorhabdus* might show presence-absence effects with presence of *Enterorhabdus* leading to decreased predictions. Lastly, high CLR-transformed abundance of genus *Family XIII AD3011 group* leads to higher predictions.

### Comparison of microbial predictors in different time-points

Independently modelling the two scenarios with varying follow-up time allowed us to compare the most relevant predictors to see if the effect and choice of microbial biomarkers remains the same. Considering metabolic outcomes that the microbiome data helped to predict, only one microbial predictor for the same metabolic outcome was shared (**Figure 3B**). Genus *UC5-1-2E3* from the *Lachnospiraceae* family was found to be among the top predictors for 2h insulin in the 18-month and 48-month time frame. Amongst the top 10 predictors for each target variable, *Escherichia-Shigella* was also shared for 2h insulin (**Supplementary Figures 1 and 2**).

The shape of the effect for *UC5-1-2E3* stays relatively stable, with extreme values for the genus showing higher predictions for both follow-up periods. This suggests that genus *UC5-1-2E3* could be considered a robust biomarker for predicting 2h insulin. Nevertheless, all other genera from the top microbial predictors were specific for a certain time frame.

## Discussion

We used machine learning to predict multiple metabolic outcomes (continuous glucose and insulin measures, HbA1c) over time periods of varying length using gut microbiome as a predictive measure. Furthermore, the modelling strategy carried out allowed us to understand the variability in performance estimates and biomarker selection. We described how high variability and personalization of the human gut microbiome leads to large variations in the performance estimates. We showed that microbial predictors can improve the prediction accuracy for continuous insulin measures and glycosylated hemoglobin, additionally highlighting differences in short and long-term cases. Finally, we identified microbial biomarkers that contribute to the improved performance and described their effect on the outcome.

Most of the current studies describing the role of bacteria in diabetes have been case-control studies with diabetes being a binary trait defined by setting a cut-off to some continuous glucose measure (3, 4, 12). Type 2 diabetes however is a disease preceded by a long-lasting prediabetic state and the development of the disease is a continuous process (13). Detailed phenotyping is definitely a strength of this study as it allows us to study the first stages of disease progression. Our results suggest that bacteria provide means for predicting changes in insulin secretion and insulin response to glucose intake. A causal effect of microbiome produced short chain fatty acids (SCFA) has been confirmed with respect to various insulin measures, primarily insulin secretion (14). **Supplementary Figure 3** shows that 2h insulin levels first increase in subjects with prediabetes, defined by the WHO classification, as a compensatory mechanism to keep glucose levels in the normal range. 2h insulin values are thus amongst the first indicators for the development of diabetes. Therefore, our results provide valuable insight into the potential application of microbiome as a predictive measure for T2D and highlight the need for detailed phenotyping in order to fully understand the role of microbiome in this disease.

Recently, Gou *et al*. (12) used a similar interpretable machine learning strategy and found bacteria that effectively differentiated type 2 diabetes cases from healthy controls in the Chinese population. Additionally, they built a microbiome risk score (MRS) and showed the causal role of identified bacteria on diabetes development after fecal microbiota transplantation to mice. The microbial predictors found do not show significant overlap with our findings. Only *Alphaproteobacteria* found by Gou *et al*. can be considered overlapping. We found one taxa from class *Alphaproteobacteria* – an uncultured genus from *Rhodospirillales* order to predict fasting insulin and 2h insulin in a 48-month time frame. We found a higher CLR-transformed abundance of unclassified *Rhodospirillales* genus decreasing type 2 diabetes risk, which is consistent with the findings of Gou *et al*. Multiple reasons might explain the observed inconsistencies. Importantly, our study was specifically designed to find prospective predictors for continuous measures. Another possible difference is the cohort structure. Our study included exclusively men, compared to 33.1% in Gou *et al*. The effect of sex on the gut microbiome is not clear, but cannot be ruled out (15, 16). Also, the metagenomic analyses of European women and Chinese subjects have shown differences which is why geographic differences in microbiome is also a possibility (3, 4).

*Rhodospirillales*, one of the strongest predictors in current study, was found to be predictive for fasting and 2h insulin in a 48-month follow-up. Order *Rhodospirillales* consists of bacteria that are known to produce acetic acid (17), which has been shown to improve insulin sensitivity (18, 19). Several other detected microbial predictors have been previously described elsewhere being associated with T2D or glucose regulation. Zhou *et al*. (20) showed that genus *Odoribacter* was negatively associated with steady-state plasma glucose which is consistent with our results for predicting 2h insulin. Krych *et al*. carried out a study on mice and identified *Muribaculaceae* (previously classified as *S24-7*) to be protective against T2D (21), which corresponds to the protective effect for HbA1c seen in our study.

Previously inconsistent associations have also been reported. We found a higher CLR-transformed abundance of *Alistipes* to predict lower values for fasting insulin, which is not consistent with the results obtained by Wu *et al*. (22), who showed positive associations with type 2 diabetes. *Subdoligranulum* has been found to be enriched in type 2 diabetes cases (23), which is inconsistent with our results as higher CLR-transformed abundance predicts lower values for 2h insulin. Similarly to the work by Gou *et al*. (12), the main reasons behind these inconsistencies are likely study design and population structure. We are not aware of any population with similar follow-up period and where microbiome data is available and oral glucose tolerance test has been carried out at the baseline and at the follow-up. Therefore, we could not replicate our findings in other populations using similar study design.

How machine learning techniques can best utilize microbiome data is still an open question (24). Therefore, the true potential of the gut microbiome for predicting T2D remains unknown. Additionally, taking the compositional nature of microbiome data into account is crucial for all types of analysis and machine learning applications (24). Previous studies have shown the advantage of using log-ratio transformations for overcoming the limitations of working with compositional data. For example, Quinn & Erb (25) and Tolosana-Delgado *et al*. (26) showed how centered log-ratio (CLR) transformed data can outperform raw proportions. Moreover, Tolosana-Delgado *et al*. (26) showed how pairwise log-ratio transformation can greatly outperform CLR transformation when a random forest algorithm is used. Thus, novel methods and strategies for handling compositionality might substantially improve the prediction accuracy for continuous metabolic outcomes.

The high variability in performance estimates shows the necessity for robust modelling strategies to achieve reliable and generalizable performance. Microbiome data are highly variable and need to be carefully analyzed. Our results show that robust model training approaches are needed for using machine learning on microbiome data. Conventionally used 10-fold cross-validation might not be sufficient to obtain generalizable models when sample sizes stay relatively small compared to the number of microbial features.

## Conclusions

In summary, our findings provide a clear indication that the microbiome can be used to predict multiple metabolic outcomes. The detailed clinical characterization and longitudinal study design of the METSIM cohort make it particularly useful for understanding host-microbiome relationships. We have identified a number of novel microbial biomarkers which could predict metabolic traits associated with pre-diabetic state. Our data provide a significant resource for further studies to determine the causal relationship of the identified biomarkers to the progression of T2D. Therefore, the prospect of using microbiome in personalized medicine is promising.

## Methods

### Study population and characterization

METSIM (Metabolic Syndrome in Men) is a randomly selected cohort of men from Eastern Finland aged 45-73 years who have been carefully phenotyped for different metabolic traits such as T2D, hypertension and obesity. We investigated a subset of the METSIM cohort that took part of the METSIM follow-up study and from whom stool samples were collected (N = 608). The data resource consists of samples taken from three time points - at baseline (baseline of METSIM 5-year follow-up study), at 18-month and at 48-month follow-up. At each time point the subjects went through a 1-day outpatient visit, during which they provided blood samples after an overnight fast and various parameters such as height, weight and blood pressure were measured and oral glucose tolerance test (OGTT) was performed. Additionally, at the baseline visit the subjects were interviewed about their history of diseases and drug usage. Full study protocol and data resources are described in Laakso *et al*. 2017 (27). All subjects have given written informed consent and the study was approved by the Ethics Committee of the University of Kuopio and was in accordance with the Helsinki Declaration.

In contrast to case-control studies, continuous “metabolic outcomes’’ (MO) were used as target variables in the modelling framework. The advantage of using continuous metabolic outcomes is that the phenotype is more distinct and there are no borderline cases with similar abilities of handling glucose as there likely are in the case-control setting (6). In total, two glucose measures, two insulin measures, glycosylated hemoglobin and three calculated glucose regulation indexes were considered (**Figure 1**). Glycosylated hemoglobin (HbA1c), fasting insulin, 2h insulin, fasting glucose and 2h glucose were measured according to the study protocol (27). Matsuda insulin sensitivity index was calculated according to (28). Insulin secretion index was calculated as Secretion index = *AUC*_*Insulin*(0–30min)_/*AUC*_*Glucose*(0–30min)_, where area under curve (AUC) was calculated using the trapezoidal formula. Disposition index was calculated as Disposition index = Secretion index ∗ Matsuda. Matsuda insulin sensitivity index and insulin secretion index have been previously shown to be best estimates for insulin sensitivity and insulin secretion in the METSIM cohort (29). Summary statistics for metabolic outcomes and additional covariates considered as predictors in the machine learning models are shown in **Supplementary Table 1**.

### Microbiome data collection, sequencing and data processing

Stool samples were collected at baseline visit during the evaluation at University of Kuopio Hospital and immediately stored at - 80°C. Microbial DNA was extracted using the PowerSoil DNA Isolation Kit (MO BIO Laboratories, Carlsbad, CA, USA) following the manufacturer’s instructions. Fecal microbiota composition was profiled by amplifying the V4 region of the 16S rRNA gene with 515F and 806R primers as previously described (30). PCR products were quantified with Quant-iTTM PicoGreen® dsDNA Assay Kit (Thermo Fisher). Samples were combined in equal amounts (∼250 ng per sample) into pools and purified with the UltraClean PCR® Clean-Up Kit (MO BIO). Sequencing was performed on an Illumina HiSeq 3000 Instrument.

Raw demultiplexed data were imported into open-source software QIIME2 (version 2019.7) using the q2-tools-import script with *CasavaOneEightSingleLanePerSampleDirFmt* input format (31). DADA2 software was used for denoising (32). DADA2 uses a quality-aware model of Illumina amplicon errors to attain an abundance distribution of sequence variance, which has a difference of a single nucleotide. *q2-dada2-denoise-single* script was used to truncate the reads at position 123, trimming was not applied. Chimera removal was done with the “consensus” filter in which chimeras are detected in each sample individually and sequences established as chimeric in a certain fraction of samples are removed. After denoising step, amplicon sequence variants (ASVs), equivalent to OTUs, were aligned using MAFFT (33) and phylogeny was constructed with the FASTTREE (34). Taxonomy assignment was done using the q2-feature-classifier with the pre-trained naïve Bayes classifier based on reference reads from SILVA 16S V3-V4 v132_99 database with similarity threshold of 99%. Seven samples didn’t pass quality control during the sequencing process and were removed from further analysis.

The average number of reads per sample was 1.351.289, samples with less than 100.000 reads were excluded from further analysis. Rest of the samples were aggregated to genus level which resulted in 553 genera. Further filtering procedure was carried out, to include only the most common genera for the prediction task. Genera that appeared in at least 50% of the samples were included in the final modelling task, 172 in total.

Due to the nature of sequencing, read counts are uninformative and must be considered relative to the total sum of reads for a given sample (35). In order to compensate for the compositional nature of the data, centered log-ratio (CLR) transformation was used as first proposed by Aitchison (36):

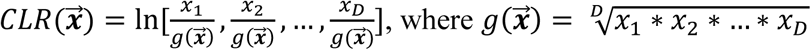

Zero replacement was carried out using R package *zCompositions* (37).

### Random forest implementation and statistical analysis

For modelling, we used samples with microbiome data available at the study baseline that did not include missing values on any of the metabolic parameters considered. In addition, subjects who had reimbursement for drug treatment of diabetes were excluded. This resulted in 529 participants for the 18-month follow-up visit and 482 participants for 48-month follow-up visit.

All random forest models were implemented in R using *caret* package and fast implementation of the random forest algorithm named *ranger* (38). Datasets were repeatedly split in 75-25 ratio to training/test datasets respectively using different seed each time. Models were tuned on training data using 10-fold cross-validation and random hyperparameter search with 100 hyperparameter combinations. Performance of the models was evaluated on the test dataset using root-mean-square error (RMSE). In case of random forest models, out-of-bag (OOB) error is also widely used to evaluate model performance. Although using out-of-bag error for evaluation can increase the sample size for model training, it has been shown that in some cases the OOB-error is largely overestimated and unreliable (39). Thus, for robust estimates, test data were used for evaluation. Permutation feature importance was used for selecting the microbial biomarkers. For explaining the obtained random forest models, accumulated local effects (ALE) plots were implemented using R package *DALEX* (40). ALE plots aim to describe the effect of a certain predictor on the metabolic outcome independently of the remaining predictors (11).

A one-tailed binomial test was carried out to test whether the probability of the model including microbial predictors outperforming the model excluding microbial predictors is greater than 0.5. Bonferroni correction was applied to assess significance (8 metabolic outcomes and two timepoints; P<0.05/16).

## Declarations

### Ethics approval and consent to participate

All subjects have given written informed consent and the study was approved by the Ethics Committee of the University of Kuopio and was in accordance with the Helsinki Declaration.

### Consent for publication

Not applicable.

### Availability of data and materials

Individual-level 16S RNA sequencing data are available in the Sequence Read Archive (SRA) under accession number PRJNA644655. All remaining phenotype data in this study are available upon request through application to the METSIM data access committee.

### Competing interest

The authors declare that they have no competing interests.

### Funding

This work was funded by Estonian Research Council grants PUT 1371 (to E.O.) and PUT1665 (to K.F.), EMBO Installation grant 3573 (to E.O.), NIH grants HL28481 (to A.J.L. and M.L.), HL144651 (to A.J.L.) and grant DK117850 (to A.J.L.). E.O. was supported by European Regional Development Fund Project No. 15-0012 GENTRANSMED and Estonian Center of Genomics/Roadmap II project No 16-0125. The METSIM study was supported by grants from the Academy of Finland (321428), Sigrid Juselius Foundation, Finnish Foundation for Cardiovascular Research, Kuopio University Hospital, and Centre of Excellence of Cardiovascular and Metabolic Diseases, supported by the Academy of Finland (to M.L.).

### Author Contributions

O.A. designed the study, performed the data analyses and wrote the manuscript. J.K and M.L designed METSIM study and oversaw collection of METSIM samples. J.L. prepared samples and performed 16S microbiome sequencing. K.L. and C.P. carried out the bioinformatic analyses from raw microbiome data. E.O. wrote and reviewed the manuscript, and supervised the data analysis. A.J.L., K.F. and M.L. reviewed the manuscript.

## Acknowledgements

We thank all METSIM study participants.

## Supplemental Material

**Supplementary Table 1.** Summary statistics for the metabolic outcomes and additional covariates included in the modelling (N = 601, seven samples were excluded in the sequencing quality control phase).

|  | Baseline | 18-months from baseline | 48-months from baseline |
| --- | --- | --- | --- |
|  | Mean (sd) | Mean (sd) | Mean (sd) |
| Age | 62.0 (5.38) | 63.6 (5.40) | 66.1 (5.36) |
| BMI | 27.8 (3.56) | 27.6 (3.63) | 27.7 (3.79) |
| HbA1c (%) | 5.6 (0.29) | 5.6 (0.27) | 5.7 (0.28) |
| Fasting glucose (mmol/l) | 5.8 (0.49) | 5.8 (0.53) | 6.0 (0.52) |
| 2h glucose (mmol/l) | 6.0 (1.99) | 5.9 (1.63) | 6.4 (1.92) |
| Fasting insulin (mU/l) | 9.5 (6.19) | 9.9 (7.12) | 10.1 (6.29) |
| 2h insulin (mU/l) | 47.8 (47.06) | 49.2 (45.29) | 55.6 (52.58) |
| Secretion index | 34.0 (20.24) | 35.6 (21.96) | 35 (20.39) |
| Matsuda index | 4.8 (3.01) | 4.7 (3.17) | 4.4 (2.95) |
| Disposition index | 125.7 (57.28) | 127.5 (67.15) | 120.9 (63.08) |
| History of elevated blood glucose | 237 (39%) |  |  |
| Diabetes in family | 222 (37%) |  |  |

**Supplementary Table 2.** Top 10 most important microbial markers for 18-month follow-up. Importance score is average permutation performance score for the variable over 200 runs. *represents taxa which were considered significant according to the average importance score.

| Trait | Phylum | Family | Genus | Average importance score |
| --- | --- | --- | --- | --- |
| 2h insulin | Euryarchaeota | Methanobacteriaceae | Methanobrevibacter | 1,64* |
|  | Firmicutes | Lachnospiraceae | [Ruminococcus] torques group | 1,46* |
|  | Firmicutes | Lachnospiraceae | UC5-1-2E3 | 1,38* |
|  | Firmicutes | Ruminococcaceae | Subdoligranulum | 1,33* |
|  | Firmicutes | Christensenellaceae | Christensenellaceae R-7 group | 1,24* |
|  | Firmicutes | Ruminococcaceae | Ruminococcaceae UCG-005 | 1,15 |
|  | Firmicutes | Lachnospiraceae | Fusicatenibacter | 1,12 |
|  | Firmicutes | Erysipelotrichaceae | Holdemania | 1,11 |
|  | Firmicutes | Peptostreptococcaceae | Terrisporobacter | 1,04 |
|  | Proteobacteria | Enterobacteriaceae | Escherichia-Shigella | 1,03 |
| HbA1c | Firmicutes | Ruminococcaceae | Ruminiclostridium 5 | 1,11* |
|  | Firmicutes | Clostridiales vadinBB60 group | uncultured bacterium | 1,07* |
|  | Bacteroidetes | Muribaculaceae | metagenome | 1,04* |
|  | Bacteroidetes | Prevotellaceae | Paraprevotella | 1,02* |
|  | Firmicutes | Clostridiales vadinBB60 group | gut metagenome | 0,99* |
|  | Bacteroidetes | Muribaculaceae | uncultured bacterium | 0,84 |
|  | Firmicutes | Clostridiales vadinBB60 group | Uncultured Thermoanaerobacterales bacterium | 0,82 |
|  | Tenericutes | uncultured organism | uncultured organism | 0,81 |
|  | Firmicutes | Clostridiales vadinBB60 group | uncultured organism | 0,78 |
|  | Firmicutes | Erysipelotrichaceae | Dielma | 0,78 |
| Secretion index | Bacteroidetes | Muribaculaceae | metagenome | 0,95* |
|  | Firmicutes | Ruminococcaceae | Papillibacter | 0,79* |
|  | Firmicutes | Ruminococcaceae | Oscillospira | 0,76* |
|  | Proteobacteria | Burkholderiaceae | Parasutterella | 0,67 |
|  | Firmicutes | Ruminococcaceae | Butyricicoccus | 0,67 |
|  | Bacteroidetes | Prevotellaceae | Alloprevotella | 0,65 |
|  | Actinobacteria | Eggerthellaceae | uncultured | 0,64 |
|  | Firmicutes | Peptococcaceae | Peptococcus | 0,63 |
|  | Firmicutes | Lachnospiraceae | Agathobacter | 0,63 |
|  | Firmicutes | Lachnospiraceae | Lachnospiraceae UCG-004 | 0,62 |

**Supplementary Table 3.** Top 10 most important microbial markers for 48-month follow-up. Importance score is average permutation performance score for the variable over 200 runs. *represents taxa which were considered significant according to the average importance score.

| Trait | phylum | family | genus | Average importance score |
| --- | --- | --- | --- | --- |
| 2h insulin | Proteobacteria | Rhodospirillales (uncultured) | gut metagenome | 1,79* |
|  | Firmicutes | Lachnospiraceae | UC5-1-2E3 | 1,73* |
|  | Firmicutes | Family XIII | Family XIII AD3011 group | 1,56* |
|  | Firmicutes | Lachnospiraceae | Shuttleworthia | 1,52* |
|  | Bacteroidetes | Marinifilaceae | Odoribacter | 1,50* |
|  | Bacteroidetes | Rikenellaceae | Alistipes | 1,46 |
|  | Firmicutes | Lachnospiraceae | CAG-56 | 1,44 |
|  | Firmicutes | Ruminococcaceae | CAG-352 | 1,44 |
|  | Proteobacteria | Enterobacteriaceae | Escherichia-Shigella | 1,42 |
|  | Firmicutes | Ruminococcaceae | Phoceia | 1,40 |
| Fasting insulin | Proteobacteria | Rhodospirillales (uncultured) | gut metagenome | 0,74* |
|  | Bacteroidetes | Prevotellaceae | uncultured | 0,65* |
|  | Bacteroidetes | Rikenellaceae | Alistipes | 0,62* |
|  | Bacteroidetes | Prevotellaceae | Prevotellaceae NK3B31 group | 0,52 |
|  | Firmicutes | Lachnospiraceae | Shuttleworthia | 0,52 |
|  | Firmicutes | Lachnospiraceae | GCA-900066575 | 0,51 |
|  | Proteobacteria | Desulfovibrionaceae | Desulfovibrio | 0,51 |
|  | Firmicutes | Christensenellaceae | Christensenellaceae R-7 group | 0,51 |
|  | Firmicutes | Christensenellaceae | uncultured | 0,49 |
|  | Bacteroidetes | Prevotellaceae | Alloprevotella | 0,49 |
| Secretion index | Actinobacteria | Eggerthellaceae | Enterorhabdus | 0,56* |
|  | Firmicutes | Erysipelotrichaceae | Asteroleplasma | 0,45* |
|  | Firmicutes | Family XIII | Family XIII AD3011 group | 0,42* |
|  | Bacteroidetes | Prevotellaceae | Prevotellaceae NK3B31 group | 0,40 |
|  | Firmicutes | Family XIII | Family XIII UCG-001 | 0,40 |
|  | Bacteroidetes | Muribaculaceae | uncultured organism | 0,40 |
|  | Firmicutes | Lachnospiraceae | [Eubacterium] xylanophilum group | 0,39 |
|  | Proteobacteria | uncultured | Azospirillum sp. 47_25 | 0,38 |
|  | Firmicutes | Ruminococcaceae | Hydrogenoanaerobacterium | 0,37 |
|  | Firmicutes | Ruminococcaceae | Ruminococcaceae UCG-010 | 0,37 |

**Supplementary Figure 1.**
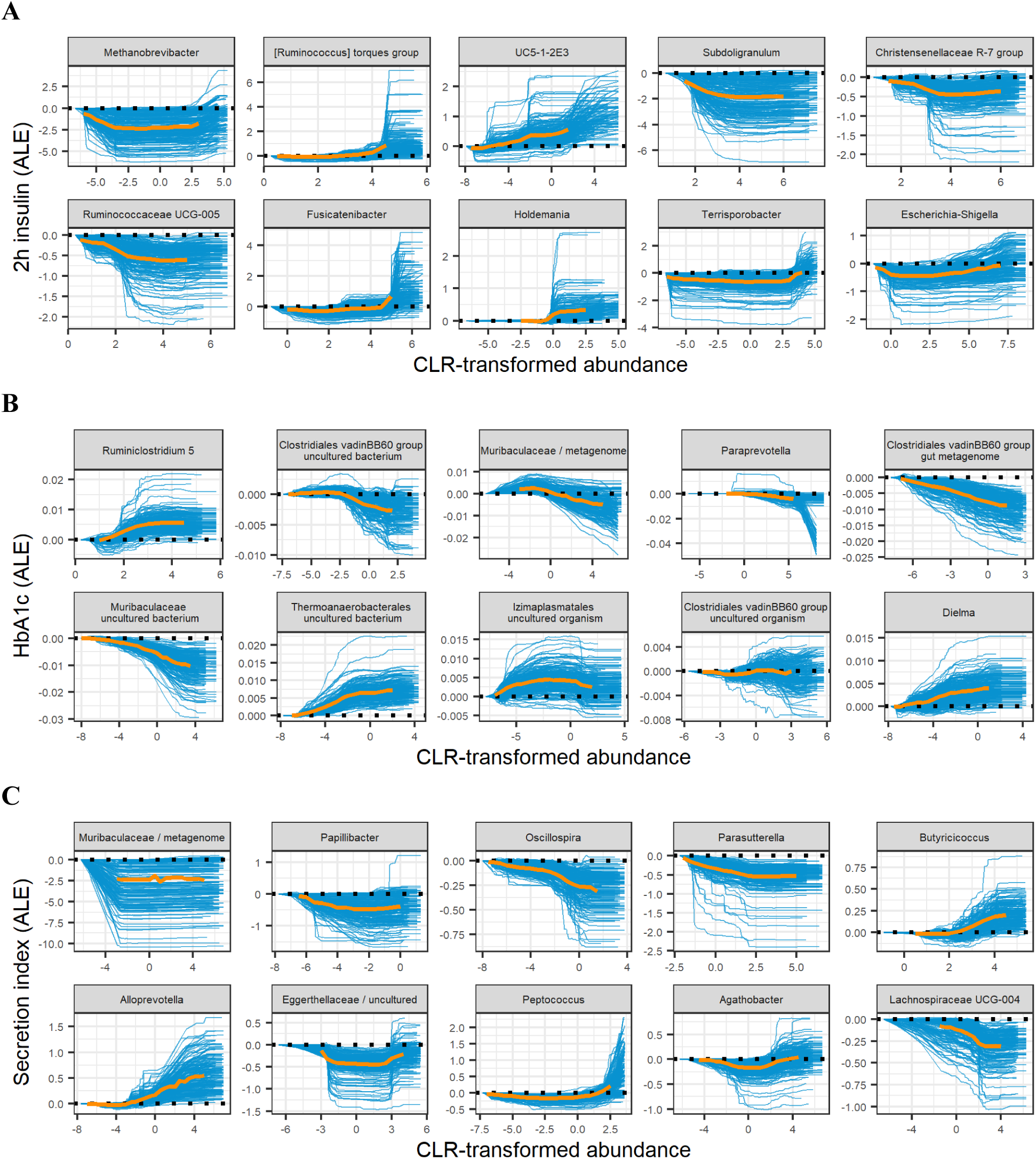
Accumulated local effect plots for the 18-month follow-up. Top 10 microbial predictors according to the average permutation importance score are displayed. Blue lines represent variable importance for each run out of 200, orange lines represent aggregated effect. Aggregated effect is displayed between the 2.5% and 97.5% quantiles of CLR-transformed abundance for the corresponding microbial marker.

**Supplementary Figure 2.**
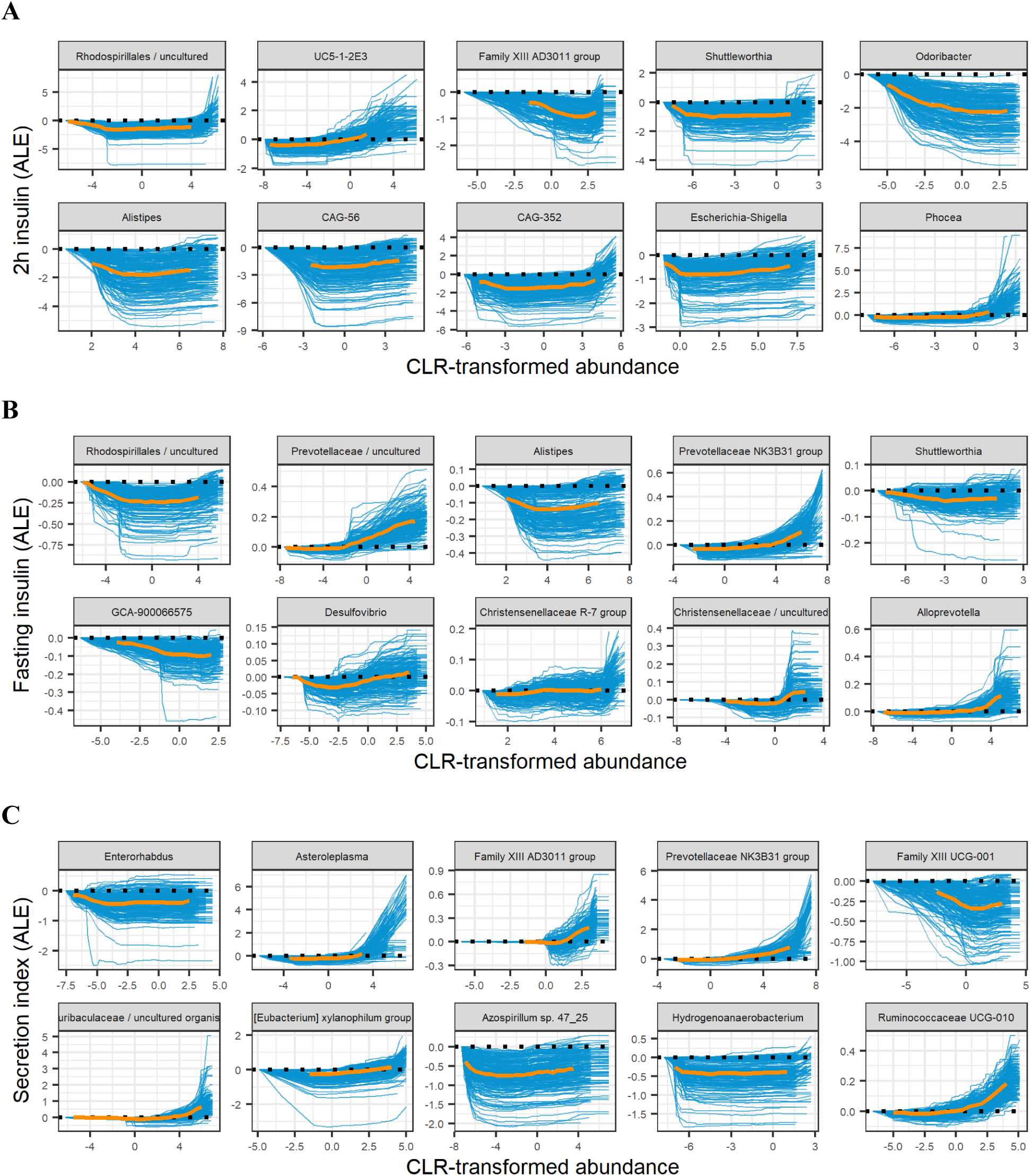
Accumulated local effect plots for the 48-month follow-up. Top 10 microbial predictors according to the average permutation importance score are displayed. Blue lines represent variable importance for each run out of 200, orange lines represent aggregated effect. Aggregated effect is displayed between the 2.5% and 97.5% quantiles of CLR-transformed abundance for the corresponding microbial marker.

**Supplementary Figure 3.**
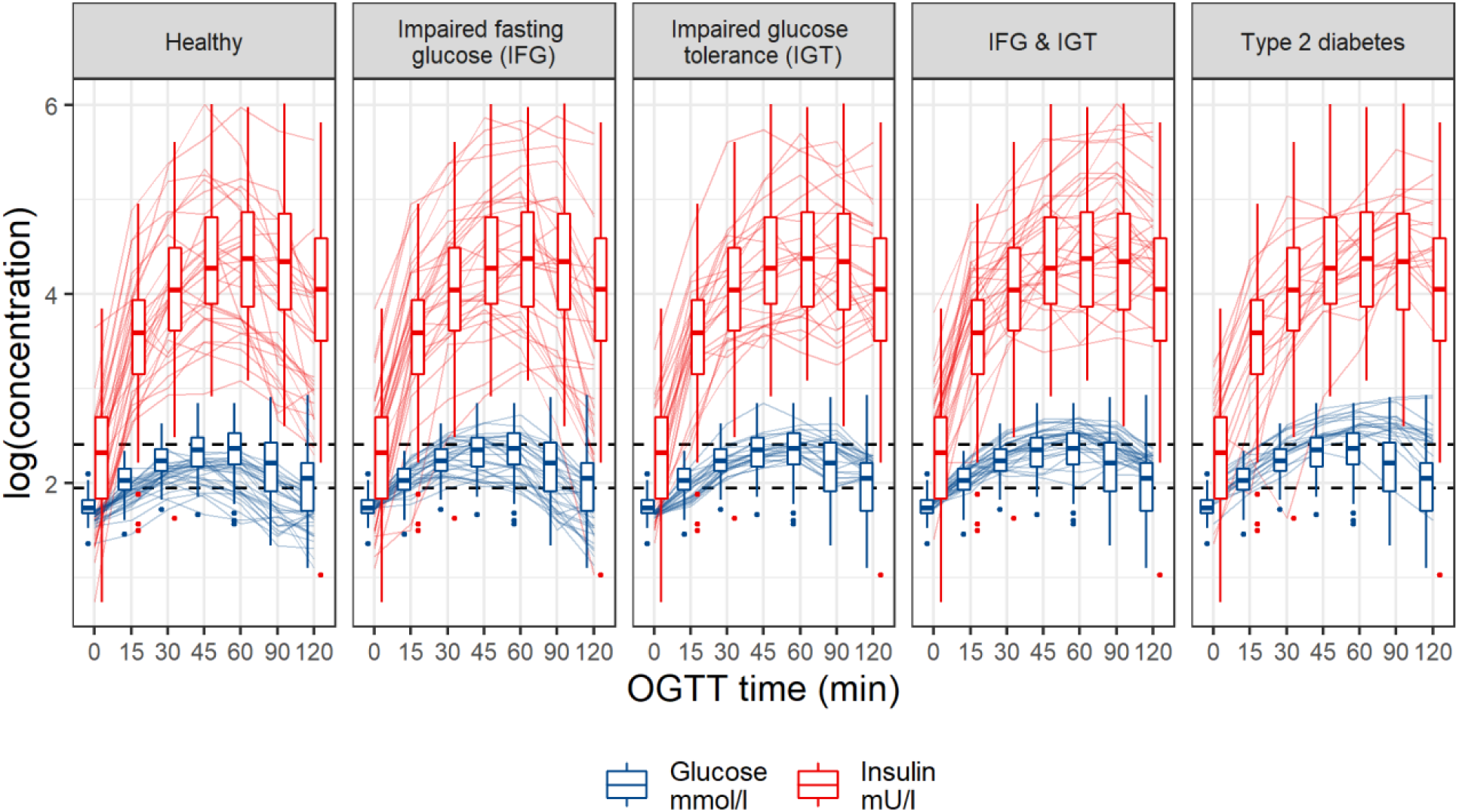
Insulin and glucose trajectories for diabetes states during oral glucose tolerance test (OGTT).

